# Evolution of host protease interactions among SARS-CoV-2 variants of concern and related coronaviruses

**DOI:** 10.1101/2022.06.16.496428

**Authors:** Edward R. Kastenhuber, Jared L. Johnson, Tomer M. Yaron, Marisa Mercadante, Lewis C. Cantley

## Abstract

Previously, we showed that coagulation factors directly cleave SARS-CoV-2 spike and promote viral entry (Kastenhuber et al., 2022). Here, we show that substitutions in the S1/S2 cleavage site observed in SARS-CoV-2 variants of concern (VOCs) exhibit divergent interactions with host proteases, including factor Xa and furin. Nafamostat remains effective to block coagulation factor-mediated cleavage of variant spike sequences. Furthermore, host protease usage has likely been a selection pressure throughout coronavirus evolution, and we observe convergence of distantly related coronaviruses to attain common host protease interactions, including coagulation factors. Interpretation of genomic surveillance of emerging SARS-CoV-2 variants and future zoonotic spillover is supported by functional characterization of recurrent emerging features.

## Introduction

The size and persistence of the viral reservoir in humans has driven considerable sequence variation among isolates of SARS-CoV-2 (Meredith et al., 2020), and distinct variant lineages have emerged (Konings et al., 2021; Kumar et al., 2021). The rapid rise and clonal expansion of the B.1.1.7 lineage (alpha variant), the B.1.617.2 lineage (delta variant), and subsequently, the B.1.1.529 lineage (omicron variant) suggest that some mutations have instilled variants with increased fitness (Harvey et al., 2021). Analysis of mutation accumulation and divergence indicates that changes in the spike S1 subunit are likely driver events in the outgrowth of emerging SARS-CoV-2 clades (Kistler et al., 2021).

The speed at which SARS-CoV-2 variants of concern have emerged has outpaced the rate at which researchers have been able to functionally characterize the effects of the mutations they harbor. The alpha, delta, and omicron variants exhibit enhanced fitness and/or escape from neutralizing antibodies, with respect to the ancestral wild type strain (Mlcochova et al., 2021; Planas et al., 2022; Shuai et al., 2022; Ulrich et al., 2022; Wang et al., 2021a). The SARS-CoV-2-S D614G substitution, which is common among VOCs, results in increased transmissibility via enhanced ACE2 binding and in hamster and ferret models (Hou et al., 2020; Korber et al., 2020; Plante et al., 2021; Zhou et al., 2021). Functional experiments have characterized the consequence of additional spike mutations on ACE2 binding (Starr et al., 2020) and escape from antibody neutralization (Chen et al., 2021; Greaney et al., 2021a; Greaney et al., 2021b; Starr et al., 2021; Wang et al., 2021b; Weisblum et al., 2020).

Coronaviruses, including SARS-CoV-2, typically require spike cleavage by host proteases at the S1/S2 boundary and S2’ site to expose the fusion peptide and enable membrane fusion and viral entry (Belouzard et al., 2009; Glowacka et al., 2011; Hoffmann et al., 2020a; Jaimes et al., 2020a; Jaimes et al., 2020c; Millet and Whittaker, 2014; Walls et al., 2020). The mechanism of cleavage activation of spike by host proteases is conserved across coronaviruses, but the cleavage recognition site is not conserved (Jaimes et al., 2020b). Viral interaction with host proteases poses a significant barrier for zoonotic spillover (Letko et al., 2020; Menachery et al., 2020) and a potential target for antiviral drugs (Hoffmann *et al*., 2020a; Hoffmann et al., 2020b). One of the most dynamic loci in the emerging lineages of SARS-CoV-2 is the S1/S2 spike cleavage site. Specifically, the P5 position, five amino acids to the N-terminal of the cleaved peptide bond, has been highly variable in the population of SARS-CoV-2. This position is subject to P681H substitution in the B.1.1.7 lineage (alpha variant) and the B.1.1.529 lineage (omicron variant); P681R substitution is present in the B.1.617.2 lineage (delta variant).

We recently discovered that coagulation factors can cleave and activate SARS-CoV-2 spike, enhancing viral entry into cells (Kastenhuber *et al*., 2022). Herein, we use FRET-based enzymatic assays to investigate the effects of mutations in SARS-CoV-2 variants of concern on interaction with factor Xa and other host proteases. Furthermore, we explored how spike cleavage sites in distantly related coronaviruses interact with various host proteases.

## Results

### Sequence divergence of SARS-CoV-2 spike codon 681 among variants of concern

Up to this point, spike codon 681, which resides in the S1/S2 cleavage site (**Fig. 1A**), is one of the highest entropy sites in the SARS-CoV-2 genome among sequenced samples (Elbe and Buckland-Merrett, 2017; Hadfield et al., 2018; Sagulenko et al., 2018). Beginning in December 2019, viral genomes have been collected globally and made available by GISAID and Nextstrain (https://nextstrain.org/), of which we visualized a subsample (Elbe and Buckland-Merrett, 2017; Hadfield *et al*., 2018). For nearly a year, SARS-CoV-2 spike encoded for proline at position 681 in almost all isolates. Samples with P681H substitution emerged in October 2020 and surpassed the frequency of P681 by March 2021 (**Fig. 1A**). Meanwhile, a P681R substitution emerged within the B.1.617.2 lineage (delta variant), and rapidly became predominant by June 2021 (**Fig. 1A**). Subsequently, the P681H substitution once again became prevalent during the clonal sweep of the Omicron variant. (**Fig. 1A**).

**Figure 1.**
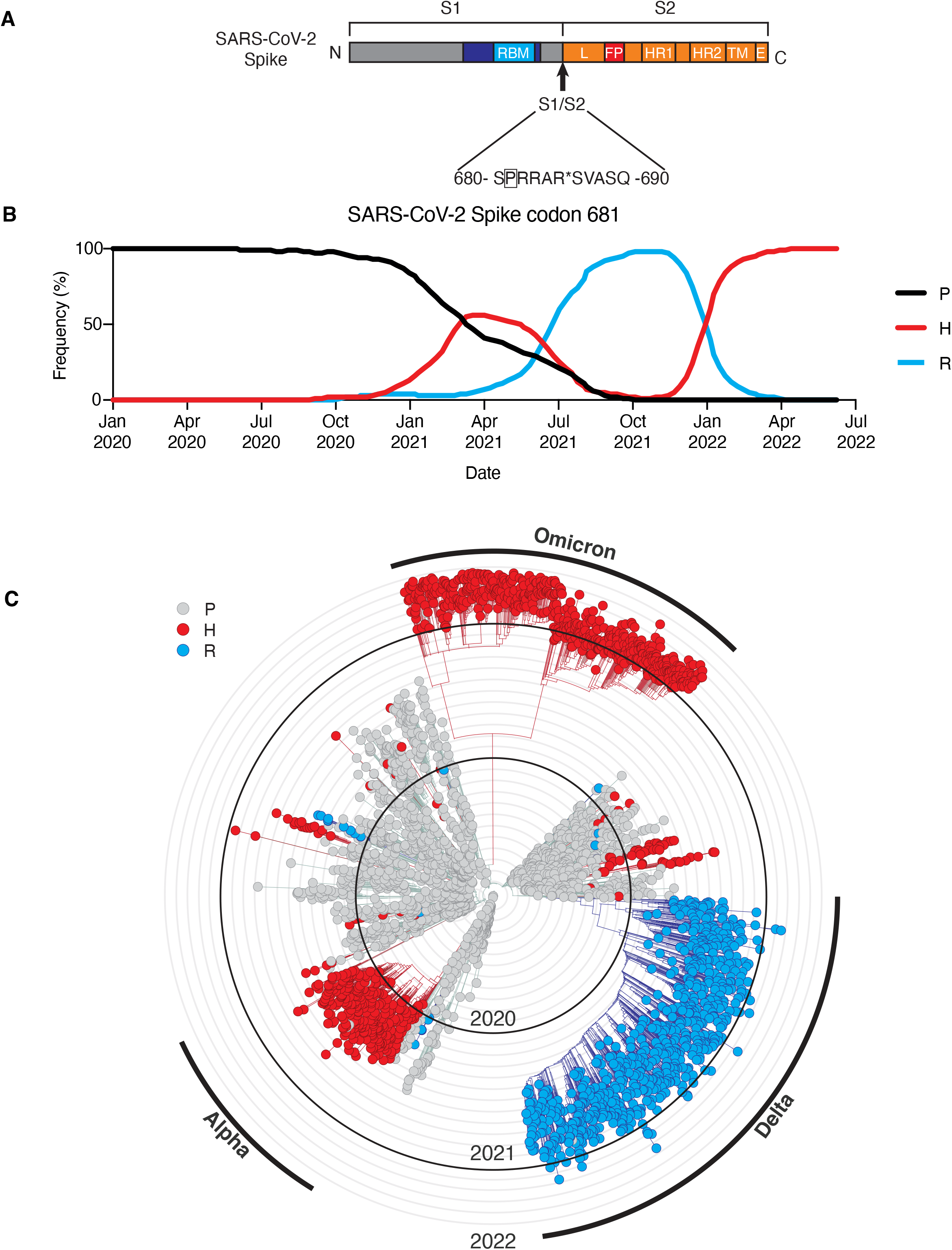
Sequence divergence of SARS-CoV-2 spike-681 among variants of concern. **(A)** Schematic of SARS-CoV-2 spike protein, highlighting position 681 adjacent to the S1/S2 site. Modified from (Kastenhuber *et al*., 2022). A subsampled collection of 3043 samples from between Dec 2109 and May 2022 from GISAID was obtained and visualized using Nextstrain (https://nextstrain.org/ncov) (Elbe and Buckland-Merrett, 2017; Hadfield *et al*., 2018). **(B)** Frequency of viral genomes sequenced with proline (black), histidine (red), or arginine (blue) at spike codon 681 by date of sample collection. **(C)** Phylogenic tree rendered by Nextstrain. Genotype at S681 of each sample is indicated by proline (gray), histidine (red), or arginine (blue). Branches corresponding to dominant variants of concern are highlighted in the outer ring.

The P681H substitution is one of many defining mutations of the B.1.1.7 lineage (alpha variant) and the P681R substitution is one of many defining mutations of the B.1.617.2 lineage (delta variant). Numerous factors may have contributed the rise in frequency of these mutations, including positive selection of other driver mutations co-occurring in the same lineage, and representation of different regions in deposited viral genomes. However, outside of the primary clades, both P681H and P681R appear to have arisen independently multiple times and shown evidence of expansion through transmission, consistent with the possibility of a functional advantage (**Fig. 1B**).

### Substitutions at SARS-CoV-2 Spike S1/S2 site cause divergent changes to interactions with host proteases

We specifically tested how substitutions observed in emerging lineages of SARS-CoV-2 variants affect cleavage of the spike S1/S2 site by various host proteases. Comparing enzyme kinetics on peptide substrates with P681 (WT, corresponding to Wuhan-Hu1) and B.1.1.7 (P681H), we found that the P681H led to an increase in factor Xa activity (**Fig. 2A**), but we found no evidence for changes in cleavability by furin, TMPRSS2, or thrombin (**Fig. 2B-D**). On the other hand, P681R substitution increased V_max_ of factor Xa by 65% as well as increasing V_max_ of furin cleavage by 99% with respect to the ancestral WT sequence (**Fig. 2A-B**). TMPRSS2 and thrombin showed decreased activity against the P681R substrate (**Fig. 2C-D**).

**Figure 2.**
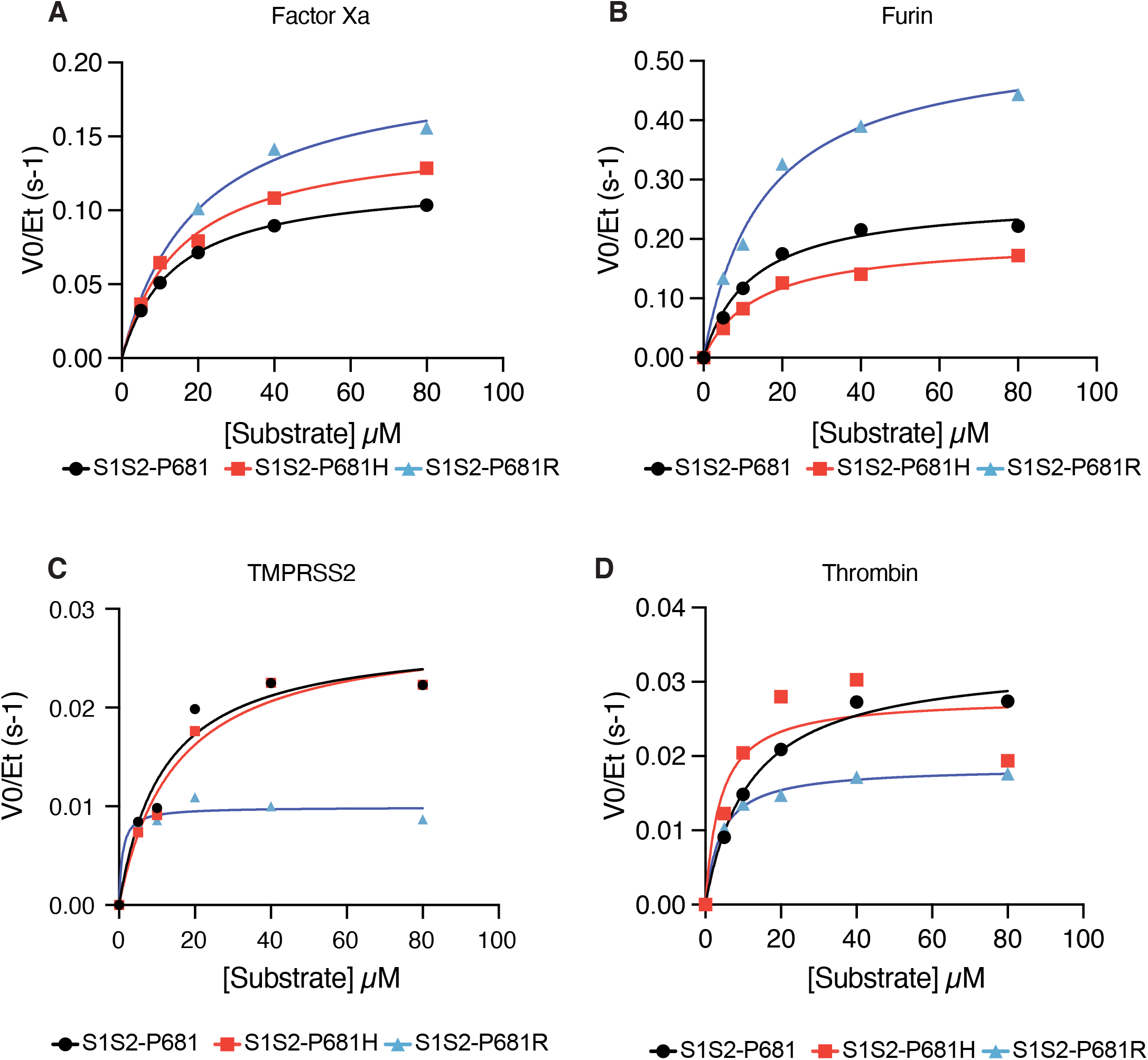
Substitutions at SARS-CoV-2 Spike S1/S2 site cause divergent changes to interactions with host proteases. Reaction rates (expressed as initial reaction velocity V_0_ normalized to the concentration of enzyme E_t_) for the cleavage of SARS-CoV-2 spike S1/S2 ancestral (P681) and variant (P681H and P681R) peptide substrates by **(A)** factor Xa, **(B)** furin, and **(C)** TMPRSS2, and **(D)** Thrombin were measured over a range of 0-80 µM substrate.

### SARS-CoV-2 spike variants remain sensitive to nafamostat

Nafamostat was found to be a multi-targeted inhibitor of TMPRSS2 as well as coagulation factors and other transmembrane serine proteases involved in viral entry (Kastenhuber *et al*., 2022). We investigated whether mutations in the S1/S2 site could affect the efficacy of nafamostat to block factor Xa-mediated spike cleavage. Although factor Xa exhibits increased V_max_ with P681H and P681R variant substrates (**Fig. 2A**), factor Xa cleavage of both variant substrates remains equivalently sensitive to nafamostat (**Fig. 3A-C**).

**Figure 3.**
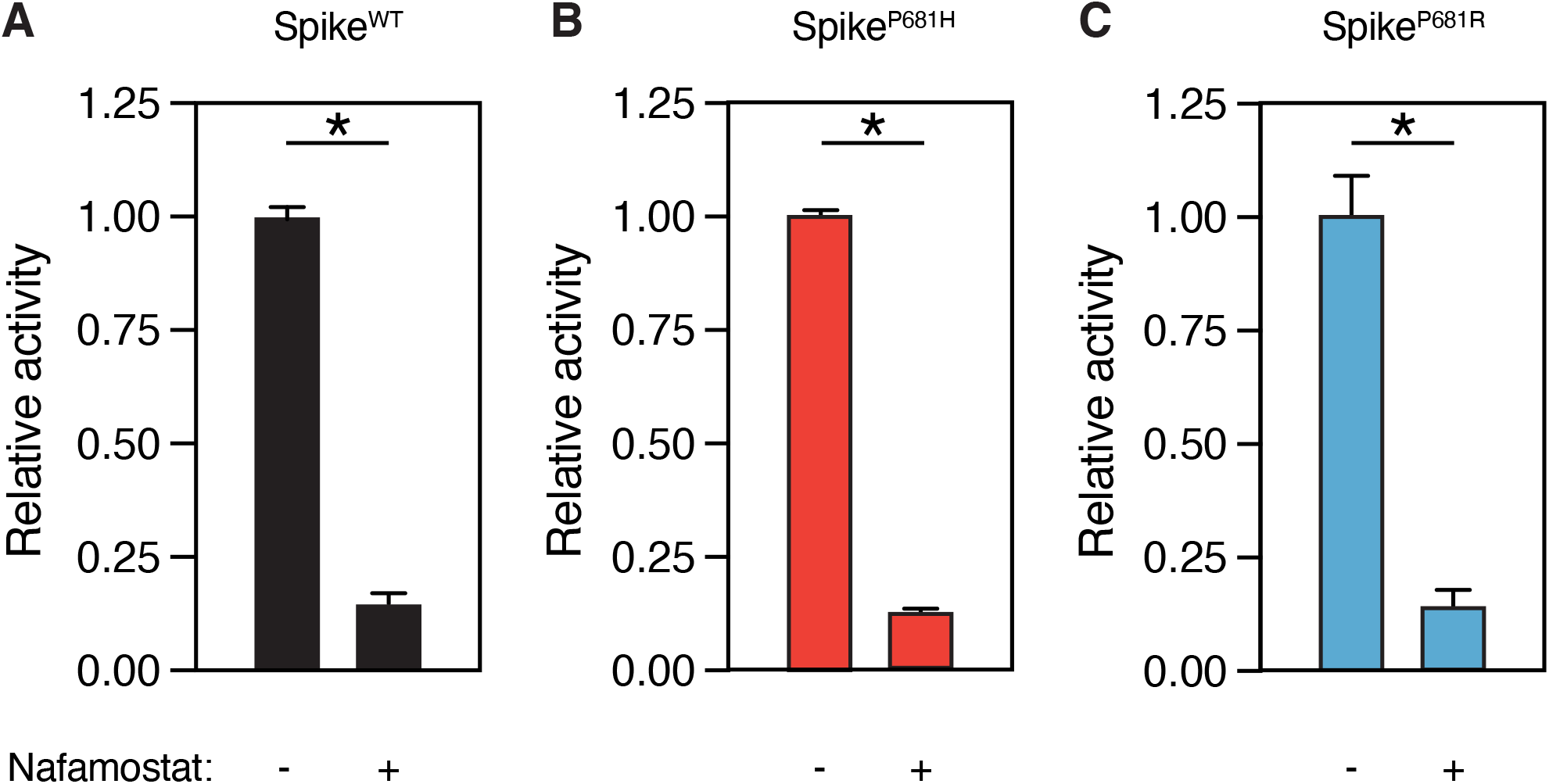
SARS-CoV-2 spike variants remain sensitive to nafamostat. Relative activity of factor Xa (125nM) with or without 10µM nafamostat in reaction with S1/S2 FRET peptide substrate (50 µM) corresponding to **(A)** WT ancestral sequence P681, **(B)** P681H substitution, and **(C)** P681R substitution. **\* P<0.05, two-tailed t-test. Error bars represent +/- SEM**.

### Effect of phosphorylation at the S1/S2 site on spike cleavage

We hypothesized that interaction with host kinases could modify interactions with host proteases. To evaluate how phosphorylation at serine residues near the S1/S2 site influence the cleavability of the site by proteases, we used singly phosphorylated peptide substrates corresponding to the S680, S686, and S689 residues **(Fig. 4A)**. Phosphorylation of Ser 680, in the P6 position upstream of the cleavage site, completely abolished furin cleavage and had a moderate impact (30-50% inhibition) on factor Xa, TMPRSS2, and thrombin cleavage **(Fig. 4B-E)**. Phosphorylation of Ser 686, in the P-1 position immediately adjacent to the cleaved amide bond, had a strong inhibitory effect on all four proteases **(Fig. 4B-E)**. Phosphorylation of Ser 689, in the P-4 position C-terminal to the cleavage site, had enzyme-specific effects on cleavage. Factor Xa and TMPRSS2 were moderately inhibited and thrombin was strongly inhibited by p-S689 **(Fig. 4B,D,E)**; however, furin cleavage was enhanced by p-S689 **(Fig. 4C)**. Post-translational modification by phosphorylation has substantial effects on the cleavability of the S1/S2 site.

**Figure 4.**
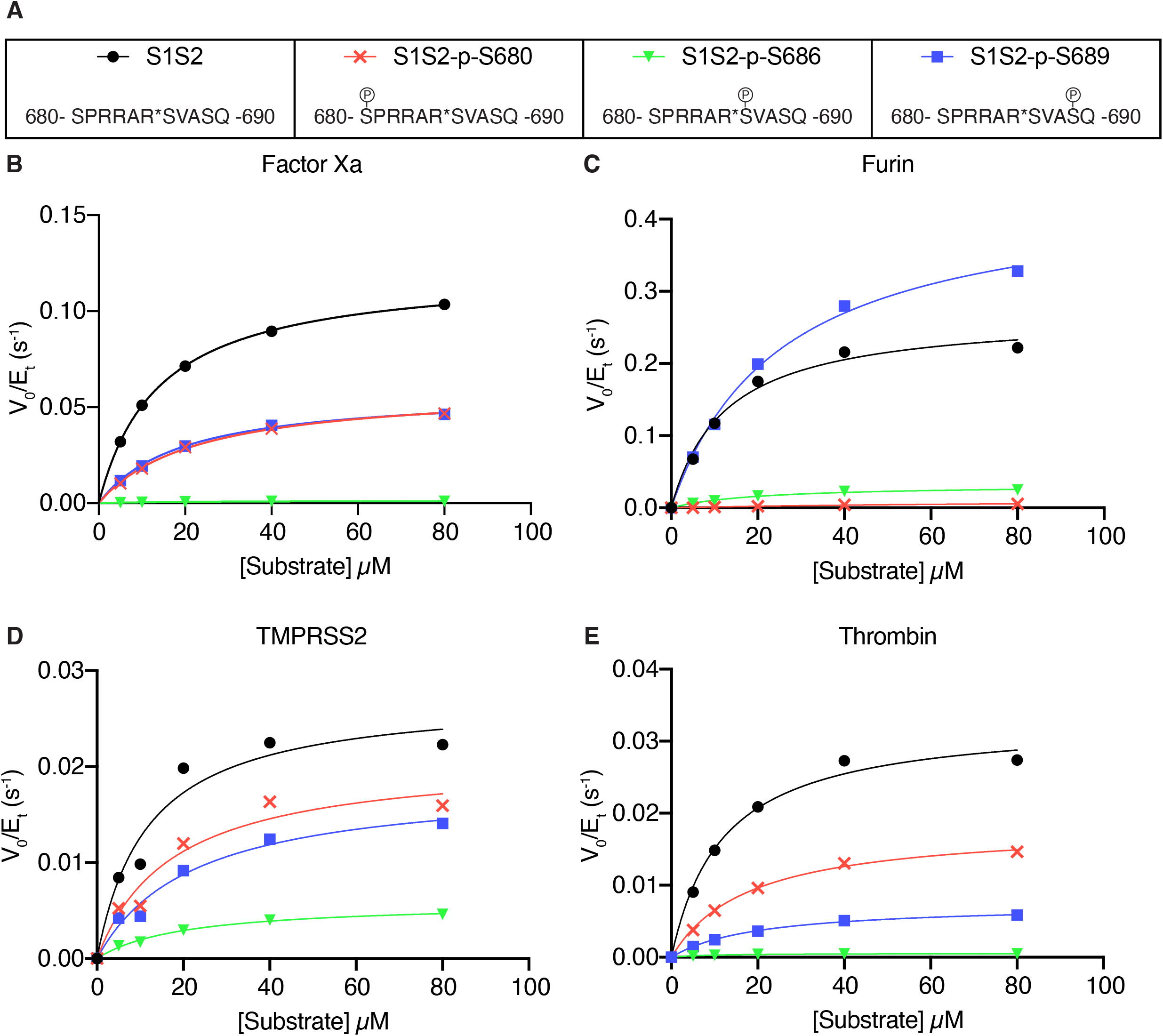
Effect of phosphorylation at the S1/S2 site on spike cleavage. **(A)** Phosphorylated peptides were generated for serine residues (S680, S686, S689). Reaction rates (expressed as initial reaction velocity V0 normalized to the concentration of enzyme Et) for the cleavage of unmodified or phosphorylated substrates by **(B)** factor Xa, **(C)** furin, and **(D)** TMPRSS2, and **(E)** Thrombin were measured over a range of 0-80 µM substrate.

**Figure 5.**
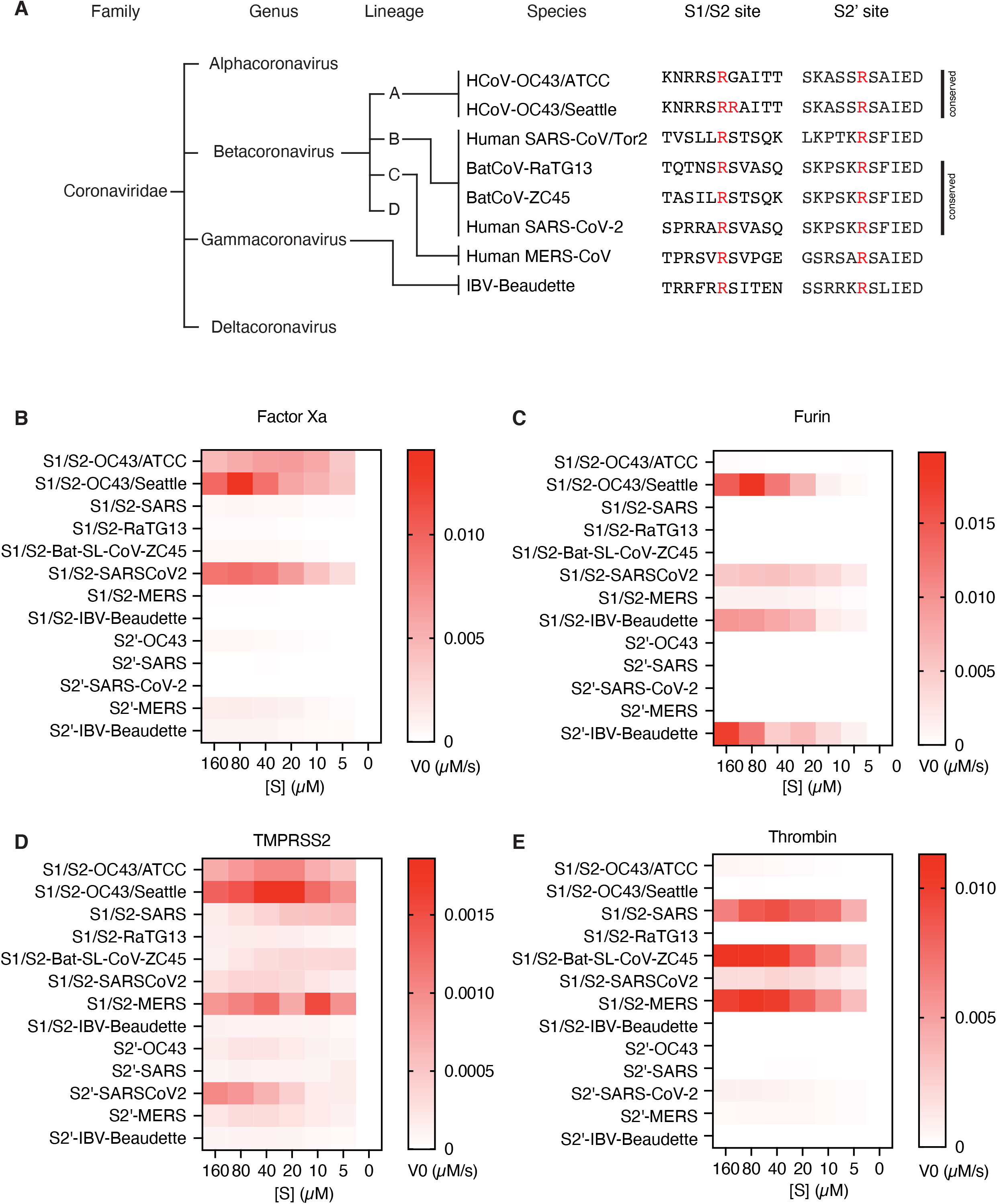
Proteolytic fingerprint of diverse coronavirus lineages. **(A)** Phylogenic relationship of a panel of coronaviruses with the corresponding aligned S1/S2 and S2’ cleavage sites. Heatmaps depicting the initial velocity V0 of cleavage of the indicated peptide substrates (rows) and concentrations (columns) by **(B)** factor Xa, **(C)** furin, **(D)** TMPRSS2, and **(E)** thrombin.

### Convergent evolution of cleavability by host proteases in diverse coronavirus species

It is not clear to what extent the cleavability by coagulation factors is specific to SARS-CoV-2 and its variants or if this is a common feature among coronaviruses. The coronaviridae family is categorized into four genera (alphacoronavirus, betacoronavirus, gammacoronavirus, and deltacoronavirus) with differences in sequence, function, and host range (Cui et al., 2019). Betacoronaviruses have evolved into four divergent lineages A-D, where lineage A contains common cold coronavirus HCoV-OC43, lineage B contains SARS and SARS-CoV-2, and lineage C contains MERS (Jaimes *et al*., 2020b) (**Fig. 4A**). We examined the interactions between host proteases and peptide substrates corresponding to a variety of betacoronaviruses from different lineages, and an outgroup avian gammacoronavirus infectious bronchitis virus (IBV-Beaudette). These substrates included diverse coronaviruses, severe and mild, zoonic and host-restricted. Interestingly, we found that no two species of coronavirus had identical susceptibility to host proteases. Only the SARS-CoV-2 S1/S2 site is cleavable by all four enzymes studied (**Fig. 4B-C, E**). In addition to SARS-CoV-2 S1/S2, factor Xa showed remarkable activity against HCoV-OC43 S1/S2 (**Fig. 4B**). A sequence from a clinical isolate of HCoV-OC43 (S1/S2-OC43/Seattle), but not the mouse-passaged laboratory strain of HCoV-OC43 (S1/S2-OC43/ATCC) was furin-sensitive (**Fig. 4B**). Furin efficiently cleaved both the S1/S2 and the S2’ sites of IBV-Beaudette, although these substates were not preferred by the other enzymes tested (**Fig. 4B**). Cleavability by thrombin was observed for the S1/S2 sites of SARS, MERS, and SARS-CoV-2, but not RatG13, a bat coronavirus with the highest known genome-wide sequence identity to SARS-CoV-2 (**Fig. 4C**). TMPRSS2 showed, on average, relatively low activity, but was active against a wider variety of both S1/S2 and S2’ substrates in the coronavirus substrate panel (**Fig. 4D**). While each coronavirus examined has a distinct set of interactions with host proteases, common solutions have been reached by distantly related viruses, suggesting convergent evolution.

## Discussion

### SARS-CoV-2 variants of concern exhibit divergent interactions with host proteases

Substitutions within the spike protease cleavage sites of SARS-CoV-2 VOCs modify viral interaction with host proteases. Spike substitution P681R increases furin cleavability, while P681H does not, in agreement with previous reports (Liu et al., 2021; Lubinski et al., 2021a; Lubinski et al., 2021b). A simplified model of SARS-CoV-2 spike activation is that furin cleaves the S1/S2 site, which potentiates either TMPRSS2 cleavage at the S2’ site or cleavage by endosomal cathepsin L at an undetermined alternative site (Bestle et al., 2020; Hoffmann *et al*., 2020a; Jaimes *et al*., 2020a; Ou et al., 2020). However, additional host proteases including other TTSPs and coagulation factors can substitute or augment these steps (Kastenhuber *et al*., 2022; Tang et al., 2021). Given that recurrent substitutions at P681 (adjacent to the S1/S2 site) have divergent effects on furin cleavage, it is likely that modified interaction with other host proteins likely contribute to selection pressure on the sequence of the S1/S2 site. For example, both P681H and P681R substitutions increase susceptibility to factor Xa-mediated cleavage. The effect of factor Xa can easily be overlooked as it is not apparent in the setting of cell culture or organoid experiments, unless added exogenously. Also, animal models of coronavirus have not been described to recapitulate coagulopathy associated with severe disease in humans (Kim et al., 2020; Leist et al., 2020; Sia et al., 2020; Zheng et al., 2021). The role of coagulation factors and other microenvironmentally-derived proteases merit further study among emerging viral variants.

### Functional characterization to support interpretation of emerging VOCs and zoonotic spillover events

The COVID-19 pandemic is an extremely challenging global health crisis, exacerbated by the continued emergence of viral variants, the impact of which can often only be seen posteriorly. Furthermore, the zoonotic spillover of SARS, MERS, and SARS-CoV-2 within the last 20 years has caused concern for additional novel coronavirus epidemics in the future. Conditions associated with heightened risk of zoonotic transmission of novel viruses include changes in the extent of human contact with wildlife and livestock, increasing urbanization and travel, and an accelerating rate of interspecies “first contacts” due to climate-induced migration (Carlson et al., 2022). Genomic surveillance is a critical tool for tracking emerging variants of SARS-CoV-2 and threats of novel species of coronavirus from other mammalian hosts (Walensky et al., 2021). However, it can be difficult to extrapolate phenotypic consequences from genomic sequence alone and fluctuations in variant prevalence can be driven by local changes in human behavior and public health policy as well as characteristics of the viral variant. The B.1.1.7 lineage (alpha variant), the B.1.617.2 lineage (delta variant), and the B.1.1.529 lineage (omicron variant) have undergone near clonal sweeps of the population of SARS-CoV-2 in humans. For unclear reasons, the P.1 (gamma variant) and B.1.526 (iota variant) lineages have faded and been displaced after their initial emergence and expansion (Annavajhala et al., 2021). Fitness advantage can be mediated by a variety of specific functional phenotypes including transmission efficiency, viral particle stability, infection cycle time, immune escape, and disease severity (Mlcochova *et al*., 2021; Wang *et al*., 2021a). The goal of functional characterization of recurrently mutated sites is to anticipate the impact of novel variants of concern and the utility of available interventions.

### Towards broad coronavirus antiviral drugs

In the first two years of the COVID-19 pandemic, vaccines and nonpharmaceutical interventions have saved many lives (McNamara et al., 2022; Mesle et al., 2021; Victora et al., 2021). Anticipating the continued evolution of SARS-CoV-2 variants and future zoonotic spillover transmission of novel coronaviruses, the development of broad-acting antivirals is an area of great interest. Coronavirus antiviral development has thus far targeted viral RdRp (Remdesevir) and viral protease Mpro (Paxlovid) (Beigel et al., 2020; Hammond et al., 2022). Host-targeted antivirals, including repurposed (Hoffmann *et al*., 2020a; Hoffmann *et al*., 2020b) and novel TMPRSS2 inhibitors (Shapira et al., 2022), have been shown to reduce viral entry. We previously demonstrated that nafamostat also inhibits both TMPRSS2 and coagulation factors, which may be a collateral benefit in anti-coronavirus activity (Kastenhuber *et al*., 2022). Although variations in the S1/S2 site sequence have resulted in enhanced factor Xa cleavability, we show here that nafamostat remains effective to block FXa-mediated cleavage of variant S1/S2 sites. Nafamostat also exhibits antiviral activity against human coronaviruses 229E and NL6, associated with milder seasonal illness (Niemeyer et al., 2021). Early, outpatient intervention with orally available drugs would be advantageous (Griffin et al., 2021), but nafamostat is an intravenous drug with a suboptimal PK profile (Quinn et al., 2022). On the other hand, intranasal delivery of nafamostat was effective in mouse models of COVID-19 and may be a promising approach (Li et al., 2021). Development of novel drugs with activity against relevant host proteases could be a valuable advancement for broad coronavirus antivirals.

### Phospho-regulation of SARS-CoV-2 spike cleavage

We found that phosphorylation of the S1/S2 site generally reduces cleavability by factor Xa, furin, TMPRSS2, and thrombin. It is understandable that a region that favors multiple basic residues for function would be inhibited by negative charge associated with phosphorylation. Phosphorylation of S680 and S686 have previously been described to inhibit furin cleavage (Ord et al., 2020). While phosphoproteomics analysis of SARS-CoV-2 viral proteins revealed numerous phosphorylation events throughout the viral proteome, no phosphorylated serine residues near the S1/S2 site have been detected (Bouhaddou et al., 2020; Davidson et al., 2020; Hekman et al., 2020; Klann et al., 2020; Stukalov et al., 2021; Yaron et al., 2020). The lack of observed phosphorylation and the robustness of SARS-CoV-2 replication would suggest that inhibitory phospho-regulation is not effective in infected cells. One might predict that selection pressure on the S1/S2 site disfavors host kinase substrate motifs so as to avoid inhibitory phosphorylation, but this does not necessarily appear to be the case (Ord *et al*., 2020)(data not shown). Alternatively, negative selection pressure through host kinase interaction could be avoided by subcellular compartmentalization of viral biogenesis, interference by other PTMs adjacent residues (including glycosylation), or exposure to host phosphatases. It is also plausible that lineage-specific expression of kinases capable of suppressing proteolytic processing of the spike could contribute to cellular tropism of SARS-CoV-2.

### Convergent evolution of host protease interactions among diverse coronavirus species

Proteolysis of coronavirus spike proteins by host proteases is clearly a selection pressure and a barrier to zoonotic spillover (Menachery *et al*., 2020). Coronavirus S1/S2 and S2’ cleavage sites exhibit distinct proteolytic fingerprints, which highlights the nuanced substrate recognition of human trypsin-like serine proteases, beyond the preference for arginine at the P1 position of the substrate (Goettig et al., 2019). The human genome encodes for more than 500 proteases and many proteases have not been sufficiently profiled to predict *in silico* which proteases are capable of cleaving a given viral sequence with any degree of certainty (Puente et al., 2005; Rawlings et al., 2018), obviating the need for direct biochemical evidence of viral interactions with host proteases.

Distantly related species of coronavirus have acquired the capacity to interact with overlapping collections of host proteases. This would suggest that selection pressure for host-mediated cleavage activation has led to convergent solutions of this critical function in multiple, independent evolutionary events. Sequence analysis has shown that furin cleavage motifs containing RXXR can be found in multiple genera of coronavirus, including a variety of betacoronaviruses (Wu and Zhao, 2020). Our data functionally confirm that furin cleavage sites, and cleavage sites of other host proteases, are widely distributed throughout coronavirus phylogeny, supporting the notion that novel protease sites emerge regularly in the evolution of coronaviruses. There has been speculation that the insertion of a polybasic sequence at the S1/S2 site of SARS-CoV-2 is suggestive of laboratory manipulation (Maxmen and Mallapaty, 2021), but this relies on the implicit assumptions that the inserted PRRA sequence has been optimized for propagation in humans and that a protease cleavage site is unlikely to emerge during natural selection. Instead, the S1/S2 site has been one of the sites in the SARS-CoV-2 genome harboring the most variation after the virus has propagated in the human population and selection for novel protease sites is a core feature of coronavirus evolution. Expanding the mechanistic depth of coronavirus host protease usage is critical to understanding coronavirus pathogenesis, to fully take advantage of genomic surveillance, and to develop pan-coronavirus antivirals.

## Author Contributions

Conceptualization, E.R.K and L.C.C.; Methodology, E.R.K, J.L.J., T.M.Y.; Investigation, E.R.K. and M.M; Writing – Original Draft, E.R.K.; Writing – Review & Editing, E.R.K. and L.C.C.; Resources, J.L.J., T.M.Y.; Funding Acquisition, L.C.C.; Supervision, L.C.C..

## Acknowledgements

The authors would like to thank Gary R. Whittaker (Cornell University), Robert E. Schwartz (WCMC), and all members of the Cantley Lab for insightful discussion and helpful comments. The efforts of the Nextstrain team and contributors to GISAID were invaluable to this study. This work was funded in part by the National Institute of Health research grant R35 CA197588 (to LCC) and the Pershing Square Foundation (L.C.C.).

## Methods

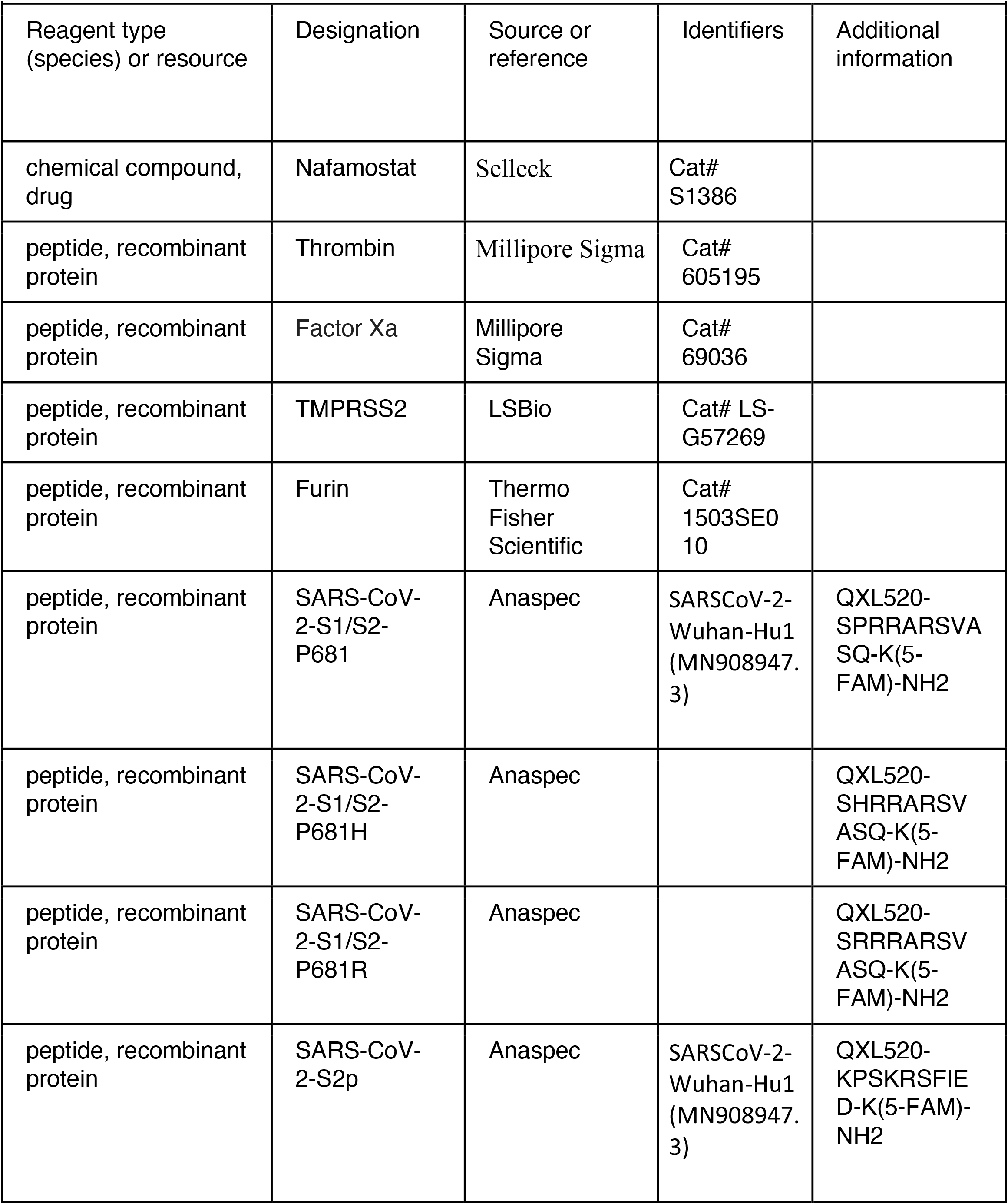
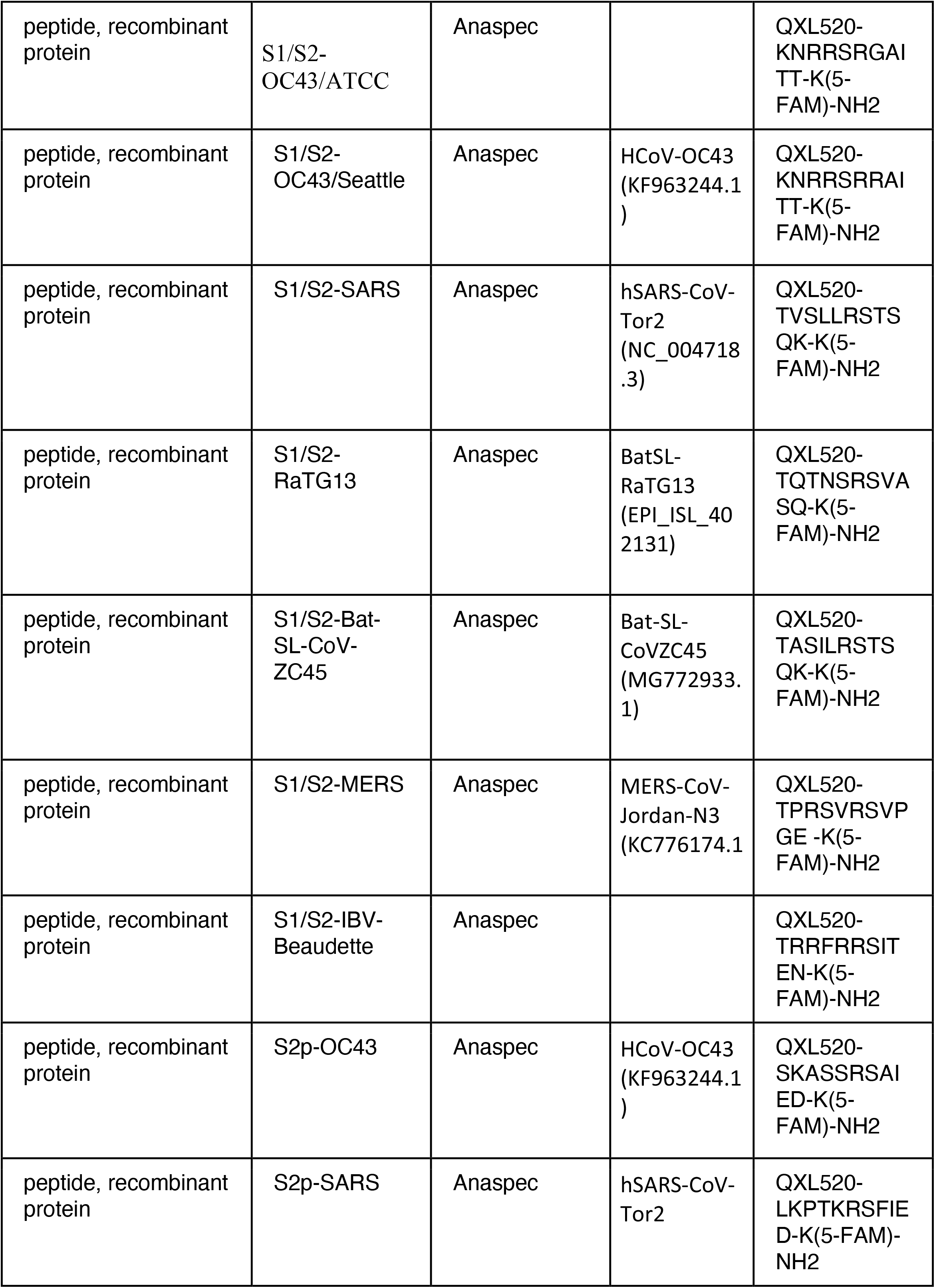
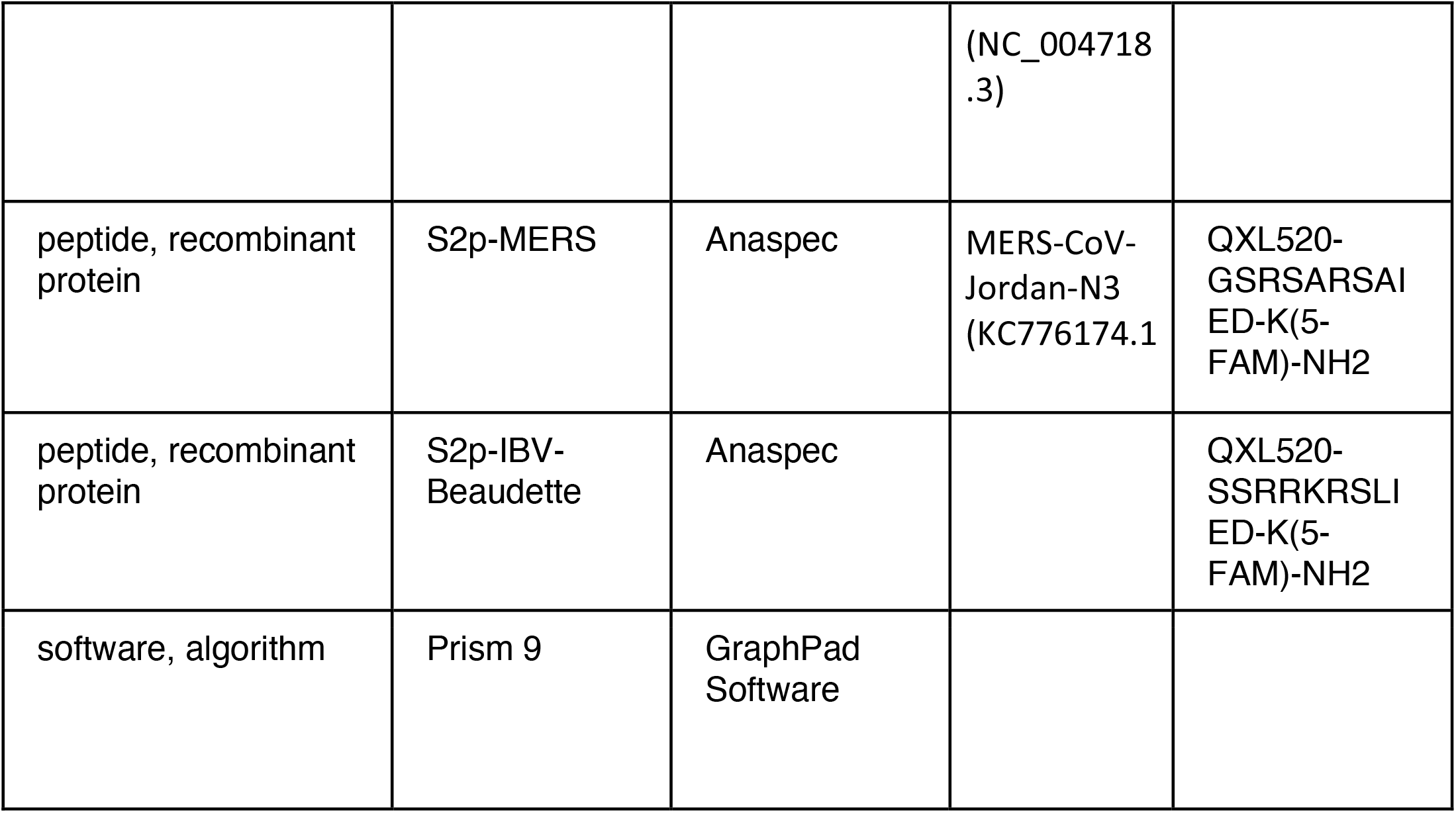
Key Resources Table.

### Sequence Analysis

A subsampled collection of 3043 samples from between Dec 2109 and May 2022 from GISAID was obtained and visualized using Nextstrain on June 3, 2022 (https://nextstrain.org/ncov/gisaid/global/all-time?c=gt-S_681&l=radial) (Elbe and Buckland-Merrett, 2017; Hadfield *et al*., 2018). Dataset parameters were set to ncov, gisaid, global, all-time. Sample clades and phylogeny were defined using default settings of Nextstrain and displayed in radial mode.

### Enzymatic Assay

Thrombin (605195) and Factor Xa, activated by Russell’s Viper Venom, were obtained from Millipore Sigma (69036). TMPRSS2, purified from yeast, was obtained from LSBio (LS-G57269). Furin was obtained from Thermo Fisher Scientific (1503SE010). FRET peptides were obtained from Anaspec and all peptide sequences are listed in the **Key resources table**. Protease assay buffer was composed of 50mM Tris-HCl, 150mM NaCl, pH 8. Enzyme dilution/storage buffer was 20mM Tris-HCl, 500mM NaCl, 2mM CaCl_2_, 50% glycerol, pH 8. Peptides were reconstituted and diluted in DMSO. Furin was used at a final concentration of 30 nM and all other enzymes were used at a final concentration of 125nM. Enzyme kinetics were assayed in black 96W plates with clear bottom and measured using a BMG Labtech FLUOstar Omega plate reader, reading fluorescence (excitation 485nm, emission 520nm) every minute for 20 cycles, followed by every 5 minutes for an additional 8 cycles. A standard curve of 5-FAM from 0-10 µM (1:2 serial dilutions) was used to convert RFU to µM of cleaved FRET peptide product. Calculation of enzyme constants was performed with Graphpad Prism software (version 9.0). Nafamostat was obtained from Selleck Chemicals.

